# Flow-sensitive K^+^ channels link flow to piezo1/PI3K/Akt1 pathway

**DOI:** 10.64898/2026.03.10.710828

**Authors:** Sang Joon Ahn, Katie Beverley, Sara T. Granados, Man Long Kwok, Jiwang Chen, Yulia Komarova, Ibra S Fancher, Shane A Phillips, Irena Levitan

## Abstract

**Background:** Endothelial response to flow is key to vascular function in health and disease. Our earlier studies demonstrated that endothelial Kir2.1 is essential for flow-induced Akt1/eNOS signaling and for flow-induced vasodilation (FIV) but the mechanistic integration between Kir and other flow signaling pathways remained poorly understood.

**Methods:** We use a combination of electrophysiological recordings in real time of flow exposure, Ca^2+^ imaging, pressure myography of resistance arteries, and echocardiography.

**Results:** We demonstrate that Kir2.1 is essential for flow-induced PI3K phosphorylation, whereas expression of myristoylated Akt1, which bypasses PI3K-dependent membrane recruitment, restores flow-induced Akt1/eNOS phosphorylation in Kir2.1-deficient endothelium. It also restores FIV in Kir2.1-deficient mesenteric arteries. We further demonstrate that Kir2.1 is essential for flow-induced Ca²⁺ influx mediated by Piezo1 and TRPV4 channels, whereas Ca²⁺ influx induced by pharmacological activation of these channels is Kir2.1 independent. Deficiency of Piezo1 does not affect endothelial Kir2.1 channels. We also discover that flow activation of endothelial Kir2.1 requires Syndecan1, thus creating a link between glycocalyx and downstream effects. Physiologically, we find that endothelial Kir2.1 is suppressed by infusion of Angiotensin-II and by advanced aging, resulting in significant impairment of FIV. In both cases, FIV is fully restored by endothelium-specific over-expression of Kir2.1.

**Conclusions:** Our study reveals that Kir2.1 serves as a mechanistic linker between endothelial glycocalyx to Piezo1-mediated Ca^2+^ influx and downstream signaling suggesting a new integrated model of endothelial mechanotransduction. A functional loss of endothelial Kir2.1 is shown to play a significant role in FIV impairment in Angiotensin-induced hypertension and aging.

## Introduction

Sensing and transducing hemodynamic forces by vascular endothelium play a pivotal role in vascular homeostasis^1,2^. One of the fundamental endothelial responses to mechanical signals is flow-induced vasodilation (FIV), endothelium-dependent vascular relaxation, an impairment of which contributes to increased blood pressure^2^. We have previously identified endothelial Kir2.1 channels as major contributors to FIV in resistance arteries^3-5^ and demonstrated that Kir2.1 are essential for flow-induced activation of eNOS via Akt signaling^4^. The mechanistic link between Kir2.1 and Akt remained, however, entirely unclear. It was also not clear what is the relationship between endothelial Kir2.1 and another mechanosensitive ion channel Piezo1, which was also proposed to constitute a primary flow sensor^6,7^. Here, we reveal new mechanistic relationships between major endothelial flow pathways identifying Kir2.1 as an essential link between flow sensing by glycocalyx to Piezo1-mediated Ca^2+^ influx and then leading to the activation of PI3K/Akt1/eNOS pathway.

Beyond acute endothelial flow signaling, we explore the role of endothelial Kir2.1 in hypertension and aging. Multiple studies showed that both hypertension and vascular aging results in endothelial dysfunction, progressive impairment of endothelial responsiveness to shear stress, the loss of the bioavailability of NO and impairment of FIV^8-10^. Our previous studies demonstrated that impairment of FIV in hypertensive patients can be partially attributed to a loss of Kir2.1-dependent vasodilation^11^. Our current study demonstrates that the loss of Kir2.1 plays a major role in Angiotensin-II (AngII)-induced and in aging-related impairment of FIV.

## Methods

Brief methods description is below and more detailed methods are provided in the Data Supplement.

### Animals and cells

All mouse experiments were approved by The University of Illinois Animal Care Committee (ACC#2022137). Mice were treated humanely minimizing pain and suffering. *<u>Primary mouse mesenteric arterial endothelial cells (MAECs)</u>* were isolated using anti-CD31 magnetic beads and maintained for up to 5 passages, as described^4^. *<u>Human adipose microvascular endothelial cells (HAMECs)</u>* were purchased from ScienCell Research Laboratories and used at low passages (P2-P4).

### Gene silencing, qPCR, Western Blot

MAECs or HAMECs were incubated with commercially designed siRNAs against RICTOR, Piezo1, Sdc1, Sdc4, or Glp1 (Qiazen) using RNAiMAX Lipofectamin and the efficiency of the downregulation was validated by qPCR. *<u>Flow-induced phosphorylation of PI3K, Akt1, eNOS and PECAM1</u>* was measured by Western Blot in cells exposed to flow using cone apparatus, as described^4^.

### Flow-induced vasodilation

First order mesenteric arteries (150-200 um) were extracted, cannulated, pressurized under 60 cmH_2_O and pre-constricted with Endothelin-1, as described^4^. Flow-induced vasodilation was determined by exposing the vessels to incremental pressure gradients with maintaining the inner pressure at 60 cmH_2_O.

### Ca^2+^ measurement

Ca^2+^ influx was measured using Fura-2 AM dye (Thermo Fisher), as described^12^. Fura-2 AM fluorescence was excited at 340 and 380 nm and collected at 510 ± 80 nm using an Axiovert 100 inverted microscope (Carl Zeiss) equipped with Plan-Apo 60× with the numerical aperture (NA) 1.4 oil immersion objective.

### Electrophysiology on freshly isolated endothelial cells

Flow-induced activation of endothelial Kir currents in freshly isolated MAECs or low-passage HAMECs were measured at 0.7 dyn/cm^2^, the half maximal activation of Kir2.1 to fluid shear^7^ using a parallel plate flow chamber with access to the recording pipette^13^. Recordings were performed using an EPC10 amplifier and accompanying acquisition and analysis software (Pulse and PulseFit, HEKA Electronik).

### Osmotic pump implantation, Echocardiogram and Blood Pressure measurement

*<u>Osmotic pump (2002, Alzet) implantation:</u>* EC-Kir2.1^WT/WT^ and EC-Kir2.1^-/-^ mice were abdominally implanted with osmotic pumps (Braintree Scientific) releasing either AngII (Cayman) or saline. *<u>Echocardiogram</u>* was recorded as described^3^. Transthoracic echocardiography was conducted using a 40 MHz transducer (MS550D) on VisualSonics’ Vevo 2100 ultrasound machine (VisualSonics, Toronto, Canada). *<u>Blood pressure</u>* was measured using CODA system (Kent Scientific)^3^. The peripheral vascular resistance is defined as the ratio of mean blood pressure (mmHg) to blood flow rate (μl/min).

### Statistics

Statistical analyses were performed using GraphPad Prizm 10. Data are presented as mean±SE. Sample size n were determined based on power analysis with 80% power and alpha 0.05. A paired or unpaired Student *t* test, 1-way or a 2-way ANOVA with or without repeated measures were used. Significance, in all cases, was set to P<0.05. Bonferroni post hoc tests were used to determine where differences existed after significance was detected with ANOVA.

## Results

### Role of Kir2.1 in PI3K/Akt1 endothelial flow signaling

We demonstrated previously that Kir2.1 deficiency in cells isolated from Kir2.1^+/-^ mice (HT) results in the loss of flow-induced phosphorylation of Akt1^3^. Here, we show that Kir2.1 deficiency abrogates flow-induced phosphorylation of PI3K, the upstream signaling event that recruits Akt to the plasma membrane^14^: Specifically, while as expected, robust flow-induced phosphorylation of PI3K is observed in MAECs isolated from WT mice, no PI3K phosphorylation is observed in MAECs isolated from Kir2.1-deficient mice. Overexpression of Kir2.1 in HT MAECs fully restores flow-induced PI3K phosphorylation (*Fig.1A*). Furthermore, genetic deficiency of Akt1 renders FIV to be insensitive to endothelial Kir2.1: no further decrease in FIV is observed when Akt1^-/-^arteries are transfected with dnKir2.1 and no rescue is observed by the over-expression of wtKir2.1 (*<u>Suppl.</u>Fig.1A<u>,B</u>*).

**Figure 1.**
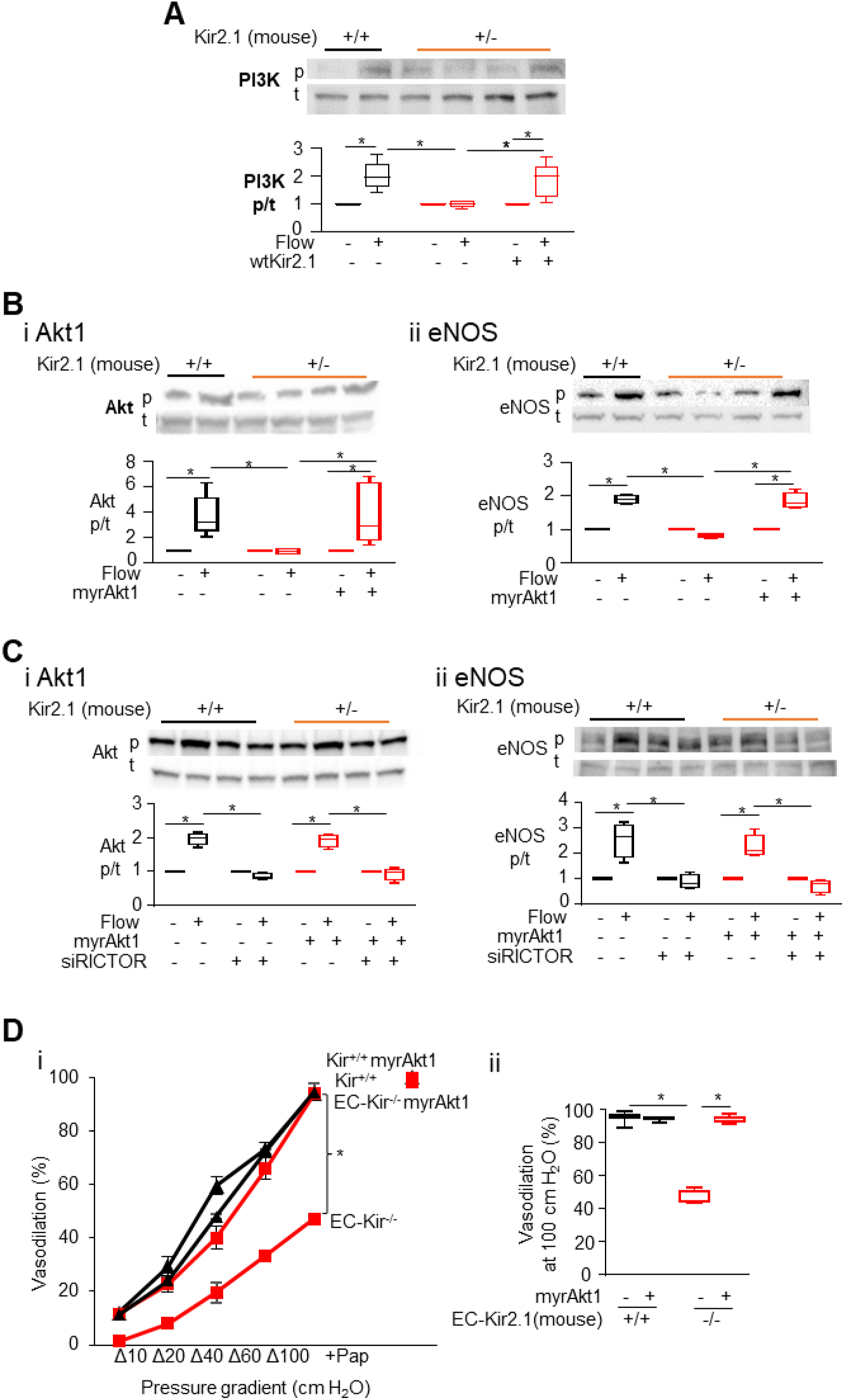
Kir2.1 regulates flow-induced Akt1 and eNOS phosphorylation via PI3K. ***<u>A</u>****<u>. Kir2.1 is required for flow-induced phosphorylation of PI3K.</u>* Representative Western blots (upper panels) and densiometric analysis (lower panels) of phosphorylated (p) and total (t) PI3K in MAECs isolated from Kir2.1^+/+^ (black) and Kir2.1^+/-^ (red) mice under static conditions or after 3 min flow of 20 dyn/cm^2^. MAECs were transfected either with empty Adv vector (labeled “–“) or with *wtKir2.1-AdvCdh5* (“+“)*. **<u>B</u>**<u>. Myr-Akt1 rescues flow-induced phosphorylation of Akt1 and eNOS in Kir2.1-deficient MAECs</u>*. Representative Western blots and densiometric analysis of Akt1 (***i***) and eNOS (***ii***) under the same flow conditions in Kir2.1^+/+^ and Kir2.1^+/-^ MAECs. The last two lanes show Kir2.1^+/-^ MAECs transfected with ***myr****Akt1-AdvCdh5* (empty Adv vector is used as control). ***<u>C</u>****<u>. Downregulation of RICTOR abrogates flow-induced Akt1 and eNOS phosphorylation</u> <u>in the presence and in the absence of myrAkt1</u>*. Representative Western blots and densiometric analysis flow-induced phosphorylation of Akt1 (***i***) and eNOS (***ii***) in MAECs transfected with empty Adv vector or *myrAkt1-AdvCdh5* with or without anti-RICTOR siRNA. *<u>A, B, C: n=5, *p<0.05</u>*. *<u>D. Endothelial expression of</u> <u>Myr-Akt1 rescues FIV in Kir2.1-deficient mesenteric arteries.</u>* FIV in mesenteric arteries from WT (black) and EC-Kir2.1^-/-^ (red) mice. Left: vasodilation curves by pressure gradient induced flow, Right: Vasodilation at 100 cmH_2_O (max flow) (n=6, *p<0.05, Pap: papaverine (100 µmol/L(µM)), endothelial independent vasodilator).

Next, we demonstrate that a mutant of Akt1, myristoylated (myr)-Akt1, which enables Akt1 translocation to the membrane independent of PI3K^15^, restores flow-induced activation of Akt1 and its downstream target eNOS in Kir2.1 deficient cells (*Fig.1Bi,ii*).

No myr-Akt1 phosphorylation is observed under static conditions, however, indicating that additional flow signaling is required. Indeed, we found that downregulating RICTOR, a key component of mTORC2^16^, abolished flow-induced phosphorylation of Akt1 and eNOS (*<u>Fig.1Ci,ii,</u> Suppl.Fig.1C*). This effect was also observed in MAECs expressing myr-Akt1 (*<u>Fig.1Ci,ii</u>*). These observations indicate that Kir2.1 is required for PI3K but not for mTORC2-dependent components of Akt1 phosphorylation.

Functionally, we demonstrate that myr-Akt1 also restores FIV in Kir2.1-deficient resistance arteries. Mesenteric arteries harvested from endothelial specific inducible Kir2.1-deficient mice (EC-Kir^-/-^) were transduced ex vivo with an adenoviral vector expressing myr-Akt1 driven by endothelial-specific promoter (Cdh5). As we have shown previously, endothelial-specific downregulation of Kir2.1 results in markedly reduced FIV compared with EC-Kir^WT/WT^ controls. Here, we demonstrate that myr-Akt1 fully restores FIV in EC-Kir^-/-^ arteries (*<u>Fig.1Di,ii</u>*).

### Role of Kir2.1 in endothelial Piezo1 signaling

Earlier studies demonstrated that a major upstream signaling step for flow-induced activation of PI3K/Akt1/eNOS pathway is activation of Piezo1 channels^17^. Our observations demonstrate that flow-induced activation of Kir2.1 channels is Piezo1 independent, but that Piezo1-mediated Ca^2+^ influx is Kir2.1 dependent. This is established first by recording Kir currents in MAECs freshly isolated from endothelial-specific Piezo1-deficient mice (EC-Piezo1^-/-^) with Piezo1^fl/fl^ used as controls. In both, EC-Piezo1^-/-^ and control cells, we observe typical Kir currents of similar amplitude that increase upon application of flow (*<u>Fig.2Ai</u>*). Flow-induced increase in Kir current density was observed in all cells ranging from 25 to 90% in individual cells (*<u>Fig.2Aii</u>*). No difference was observed in flow-sensitivity between EC-Piezo1^-/-^ and control cells (*<u>Fig.2Aiii</u>*), indicating that Kir2.1 function is independent of Piezo1.

In contrast, we found that Kir2.1 deficiency virtually abrogates flow-induced Ca^2+^ increase (*Fig.2B, Suppl.Fig.2B*), known to be mediated by Piezo1 and TRPV4 channels^6,12,18,19^. As expected, WT cells develop an immediate robust transient Ca^2+^ response to the application of flow (20 dyn/cm^2^), demonstrated by Fura2 imaging (*<u>Fig.2Bi</u>*, upper panels, *<u>Fig.2Bii</u>*). However, flow-induced Ca^2+^ response in Kir2.1-deficient cells was abolished (*<u>Fig.2Bi</u>*, middle panels, *<u>Fig.2Bii</u>*). To verify that flow-mediated Ca^2+^ response in MAECs is mediated by Piezo1, cells were transfected with Piezo1 siRNA, which eliminated >90% of flow-induced Ca^2+^ response (*<u>Fig.2Bi,</u>* lower panels*<u>, Fig.2Bii,</u> Suppl.Fig.2A*). In addition, we tested the contribution of TRPV4 channels, which are also implicated in endothelial flow-induced Ca^2+^ response^18^. Inhibition of TRPV4 with 100 nmol/L GSK219, a specific TRPV4 inhibitor^20^, resulted in a ∼60% decrease in flow-induced Ca^2+^ response. Combined downregulation of Piezo1 and inhibition of TRPV4 completely abolished the Ca^2+^ response (*Suppl.Fig.2B*).

**Figure 2.**
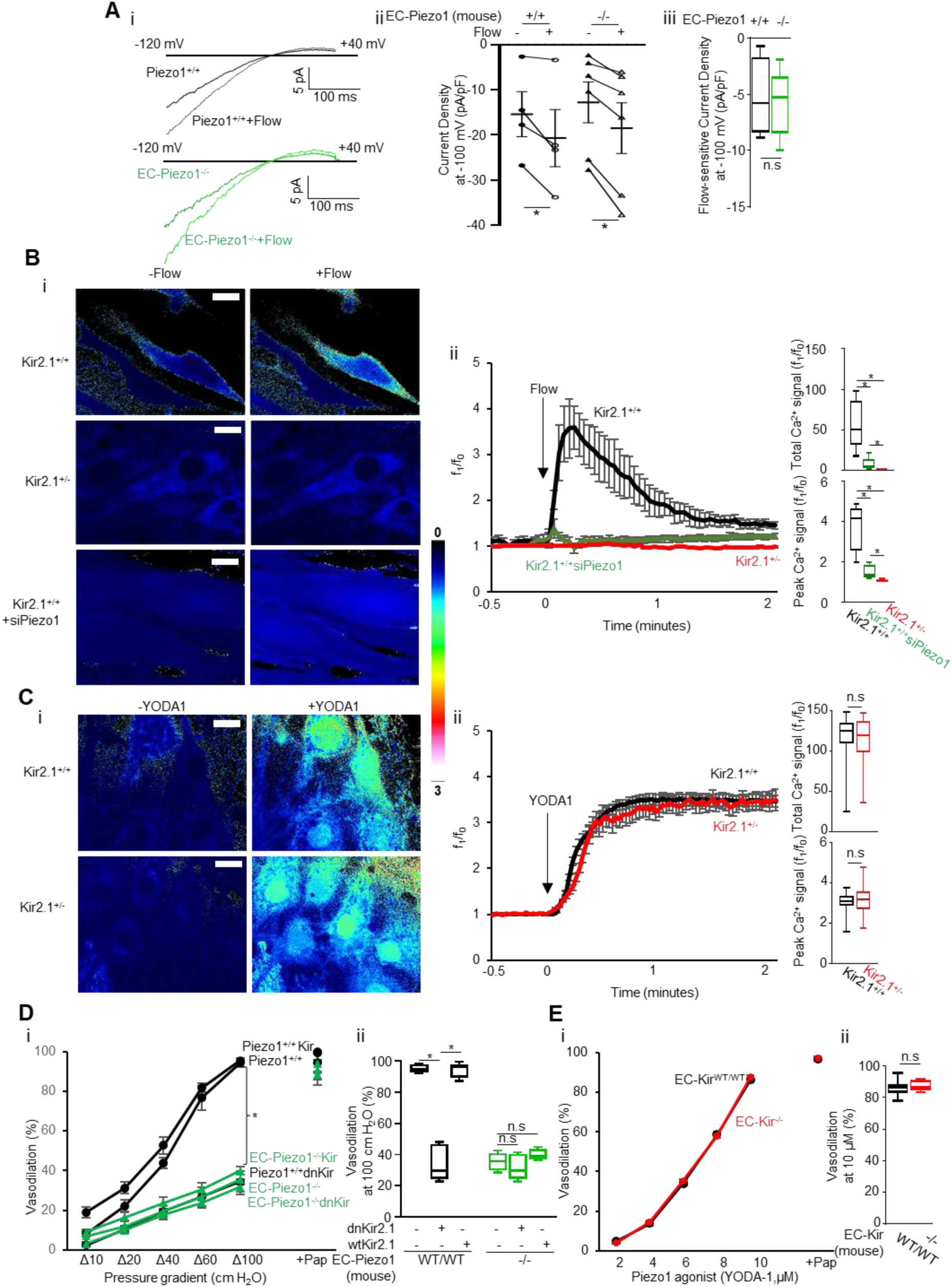
Flow activation of endothelial Kir2.1 is upstream of Piezo1-mediated Ca^2+^ influx. <u>A. Flow activation of Kir2.1 is intact in the absence of Piezo1.</u> i. Representative flow-induced Kir2.1 currents in freshly isolated MAECs from WT and EC-Piezo1^-/-^ mice under static and flow conditions. *ii*. Kir2.1 current densities under static and flow conditions at -100 mV in individual WT or EC-Piezo1^-/-^ cells (*p<0.05) *iii*. Flow-sensitive current density (flow-static). *<u>B. Kir2.1 deficiency abrogates flow-induced</u> Ca^2+^ influx*. *i.* Representative Fura-2 images of Ca^2+^ influx with and without flow (20 dyn/cm^2^) in Kir2.1^+/+^, Kir2.1^+/-^ and Kir2.1^+/+^ transfected with anti-Piezo1 siRNA (siPiezo1) MAECs (*bar: 5 µm*). *ii*. Time-dependent Ca^2+^ influx curves under flow (left), and quantified total and peak signals (right). (Kir2.1^+/+^: black, Kir2.1^+/-^: red, Kir2.1^+/+^+siPiezo1: green, in n=5 independent experiments, *p<0.01). *<u>C. Kir2.1</u> deficiency has no effect on Yoda-induced Ca^2+^ influx*. *i.* Representative images of Ca^2+^ influx with and without YODA1 (10 µmol/L(µM) (*bar: 5 µm*). *ii.* Ca^2+^ influx curves (left), and quantified total and peak signals (right). *<u>D. Functional downregulation of Kir2.1 and Piezo1 deficiency have similar non-cumulative inhibitory effects on FIV.</u> i.* FIV curves in arteries from EC-Piezo1^+/+^ (black) and EC-Peizo1^-/-^ (green) mice transfected with empty Adv vector, *wtKir2.1-AdvCdh5 (Kir),* or *dnKir2.1-AdvCdh5 (dnKir)*. *ii*. FIV at Δ100 cm H_2_O (max flow) (n=6, *p<0.05). *<u>E. Downregulation of Kir2.1 has no effect on Yoda1-induced vasodilation</u>*. *i*. YODA1-induced vasodilation curves in arteries from EC-Kir2.1^+/+^ (black) and EC-Kir2.1^-/-^ (red). *ii.* Vasodilation at 10 µmol/L(µM). (n=6) (Pap: papaverine (100 µmol/L(µM))

Our data also indicate that the requirement for Kir2.1 is specific for flow-induced Ca^2+^ response, whereas it has no effect on pharmacological activation of either Piezo1 or TRPV4. Specifically, we show that exposure to the Piezo1-specific activator Yoda1 (10 µmol/L) elicits the same Ca^2+^ response in WT and Kir2.1-deficient MAECs (*<u>Fig.2Ci,ii</u>*). Similarly, Kir2.1 expression did not alter Ca²⁺ increase in response to TRPV4 activator, GSK101 (*Suppl.Fig.2C*).

Further evidence for Kir2.1 and Piezo1 contributing to endothelial flow response via the same pathway comes from analyzing their contribution to FIV. Endothelial-specific genetic deletion of Piezo1 or endothelial-specific downregulation of Kir2.1 with dnKir2.1 decrease the FIV response to the same degree (∼50%), with no additional decrease when the two conditions are combined (*<u>Fig.2Di,ii</u>*). Also, similarly to Akt1^-/-^ arteries, FIV in EC-Piezo1^-/-^ arteries is not restored by Kir2.1 over-expression of Kir2.1. However, Yoda1-induced vasodilation is not affected: the same vasodilatory dose response to Yoda1 is observed in EC-Kir^WT/WT^ and EC-Kir⁻^/⁻^ arteries (*<u>Fig.2Ei,ii</u>*). Similarly, Kir2.1 plays no role in TRPV4-mediated vasodilatory response induced by a TRPV4 pharmacological activator, GSK101 (*Suppl.Fig.2D*). Moreover, blocking TRPV4 with GSK219 decreases FIV in control but not in EC-Kir^-/-^ arteries (*Suppl.Fig.2E*). These findings indicate again that the role of Kir2.1 in Piezo1-mediated responses is specific to flow.

Next, we tested the impact of Kir2.1 deletion on FIV while blocking P2Y₂/_Gαq/11_/PLCβ cascade, which is known to connect Piezo1 to the activation of PECAM1–VE-cadherin–VEGFR2^17^. Pharmacological inhibitors targeting P2Y₂ (AR-C118925XX), _Gαq/11_ (YM254890), or PLCβ (U73122) significantly reduced FIV in EC-Kir^WT/WT^ arteries, but none produced additional suppression in EC-Kir^⁻/⁻^ arteries (*Suppl.Fig.3A,B*). Furthermore, flow-induced phosphorylation of PECAM1 was markedly reduced in Kir2.1-deficient MAECs compared with WT controls (*Suppl.Fig.3C*).

### Syndecan-1 is required for flow-induced Kir2.1 activation

The findings described above identify Kr2.1 as an essential step in flow-induced transduction upstream of Piezo1 and TRPV4 channels. Here, we determine the upstream flow sensor required for activating Kir2.1. Our earlier study demonstrated that enzymatic degradation of endothelial glycocalyx heparan sulphate (HS) abrogates the flow sensitivity of Kir2.1^5^. To identify the HS glycocalyx element required for the flow activation of Kir2.1, three major core proteins/proteoglycans of endothelial HS glycocalyx, Syndecan-1 (Sdc1), Syndecan-4 (Sdc4), and Glypican-1 (Gpc1), Sdc1, Sdc4, or Gpc1 were silenced using siRNA (mRNA levels were reduced by more than 90% for each, *Fig.3A*). We found that Kir2.1 are sensitive specifically to SDC1: downregulation of Sdc1 abrogated Kir2.1 response to flow (*Fig.3B<u>,C</u>*), while downregulation of Sdc4 or Gpc1 had no or little effect on Kir flow response (*Fig.3C*). These observations identify the specific glycocalyx element that transduces the flow signal to Kir2.1.

**Figure 3.**
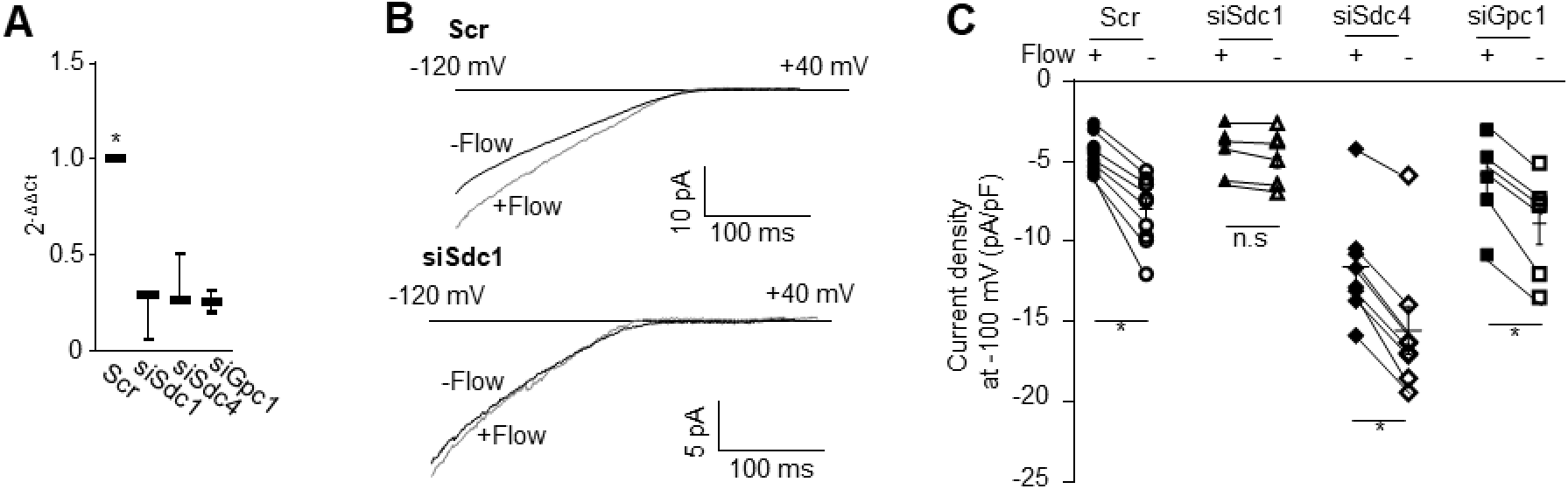
Flow-induced activation of endothelial Kir2.1 requires Syndecan1. <u>A.</u> *<u>Efficiency of mRNA downregulation for Sdc1, Sdc4, and Gpc1</u>* in HMAECs with siRNA against each gene (n=3, *p<0.05). *<u>B. Flow-induced activation of Kir2.1 is abrogated by siSDC1</u>*. Representative Kir2.1 currents under static and flow conditions in HMAECs transfected with scrambled siRNA (upper) and siSdc1 (lower). *<u>C. Kir2.1 current densities in HMAECs transfected siSdc1, siSdc4 or siGpc1</u>* or scrambled controls (n=5, *p<0.05).

### Loss of endothelial Kir2.1 causes elevation of blood pressure and peripheral resistance

We also reported previously that the global Kir2.1 deficiency elevates blood pressure and increases peripheral resistance^3^. In this study, we demonstrate that endothelial-specific deletion of Kir2.1 is sufficient to elevate systolic, diastolic, and mean arterial blood pressures (*Fig.4A*), as well as result in a significant rise in peripheral vascular resistance (*Fig.4A<u>, right most</u>*). Kir2.1 deficiency had same effects on blood pressure in male and female mice (*Fig.4A*, *Suppl.Fig.4A,B,C,D*). No significant effect of endothelial Kir2.1 deficiency was observed across an array of cardiac function parameters, including cardiac output (*<u>Fig.4Bi,ii</u>*), LV mass, ejection fraction, and fraction shortening (*Suppl.Fig.4E,F,G*). Since there was no difference in the effects of Kir2.1 deficiency on blood pressure between males and females, the cardiac parameters were measured in male mice only.

**Figure 4.**
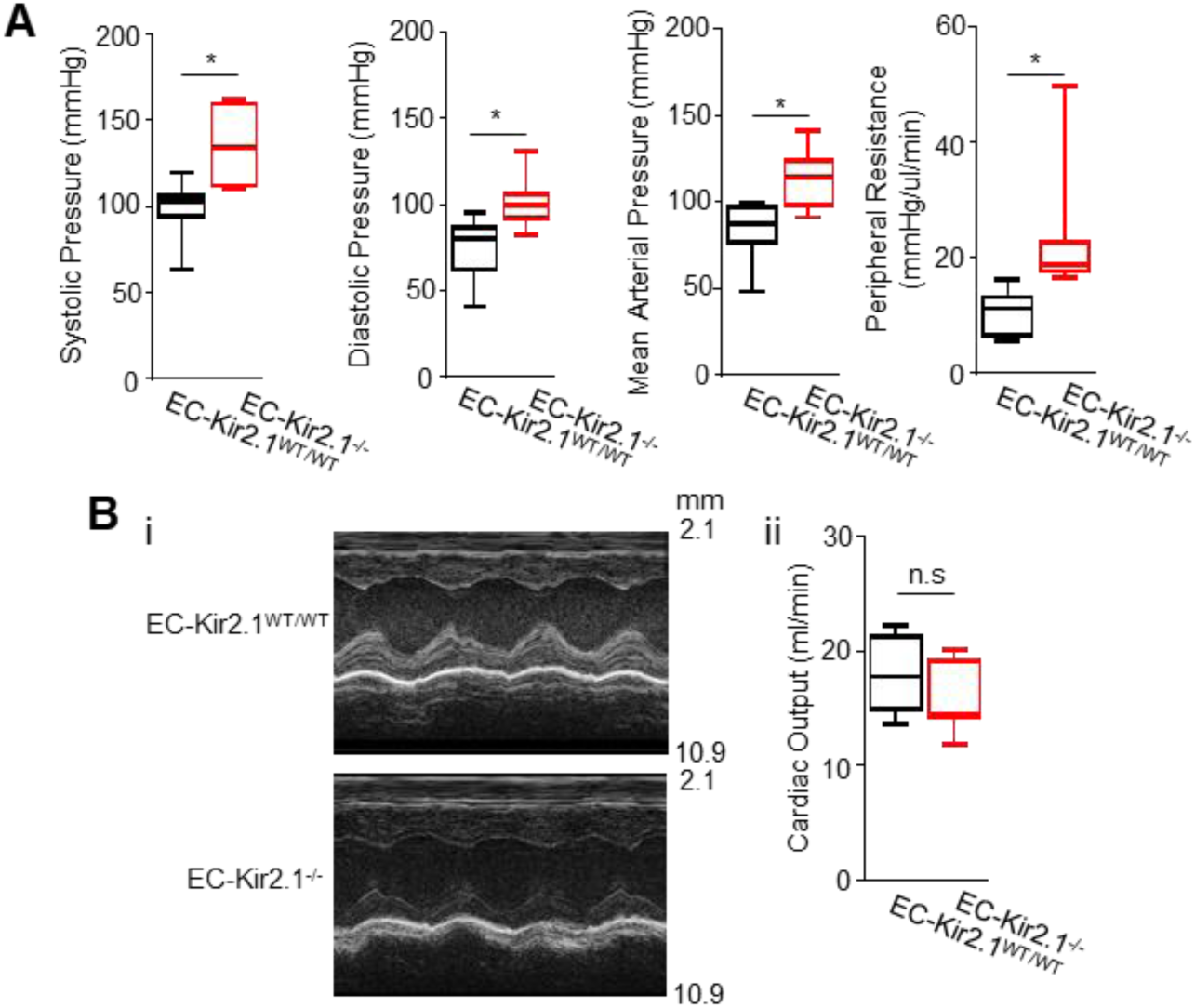
Endothelial Kir2.1 regulates blood pressure and peripheral resistance. *<u>A. Systolic, diastolic, mean arterial pressure, and peripheral resistance</u>* measured with tail-cuff in EC-Kir2.1^+/+^ (black) and EC-Kir2.1^-/-^ (red) mice (means ± SEM, n=8, *p<0.05). *<u>Endothelial Kir2.1 deficiency has no effect on cardiac output</u>. i.* Representative M-mode echocardiogram of the left ventricle of EC-Kir2.1^+/+^ and EC-Kir2.1^-/-^ mice. ii. Mean cardiac outputs of EC-Kir2.1^+/+^ (black) and EC-Kir2.1^-/-^ (red) mice (means ± SEM, n=8).

### AngII-induced impairment of FIV is mediated by the loss of Kir2.1 current

Earlier studies showed that infusion of AngII in mice causes elevation of the blood pressure and impairment of FIV^10^. Here, we demonstrate that endothelial Kir2.1 plays a key role in AngII-induced FIV impairment.

As expected, a significant increase in blood pressure observed after 7 days of AngII infusion (3 mg/kg/day) and maintained for the duration of the experiment (14 days) (*Fig.5A*). Also, consistent with the data presented above, EC-Kir^-/-^ have increased blood pressure prior to the application of AngII (day 0). Here, we demonstrate that AngII infusion suppresses endothelial Kir2.1 currents and Kir2.1-dependent component of FIV: MAECs isolated from mice infused with AngII had a significant (4-fold) reduction of Kir currents compared to control cells isolated from mice infused with saline (*Fig. 5Bi<u>,ii</u>*).

**Figure 5.**
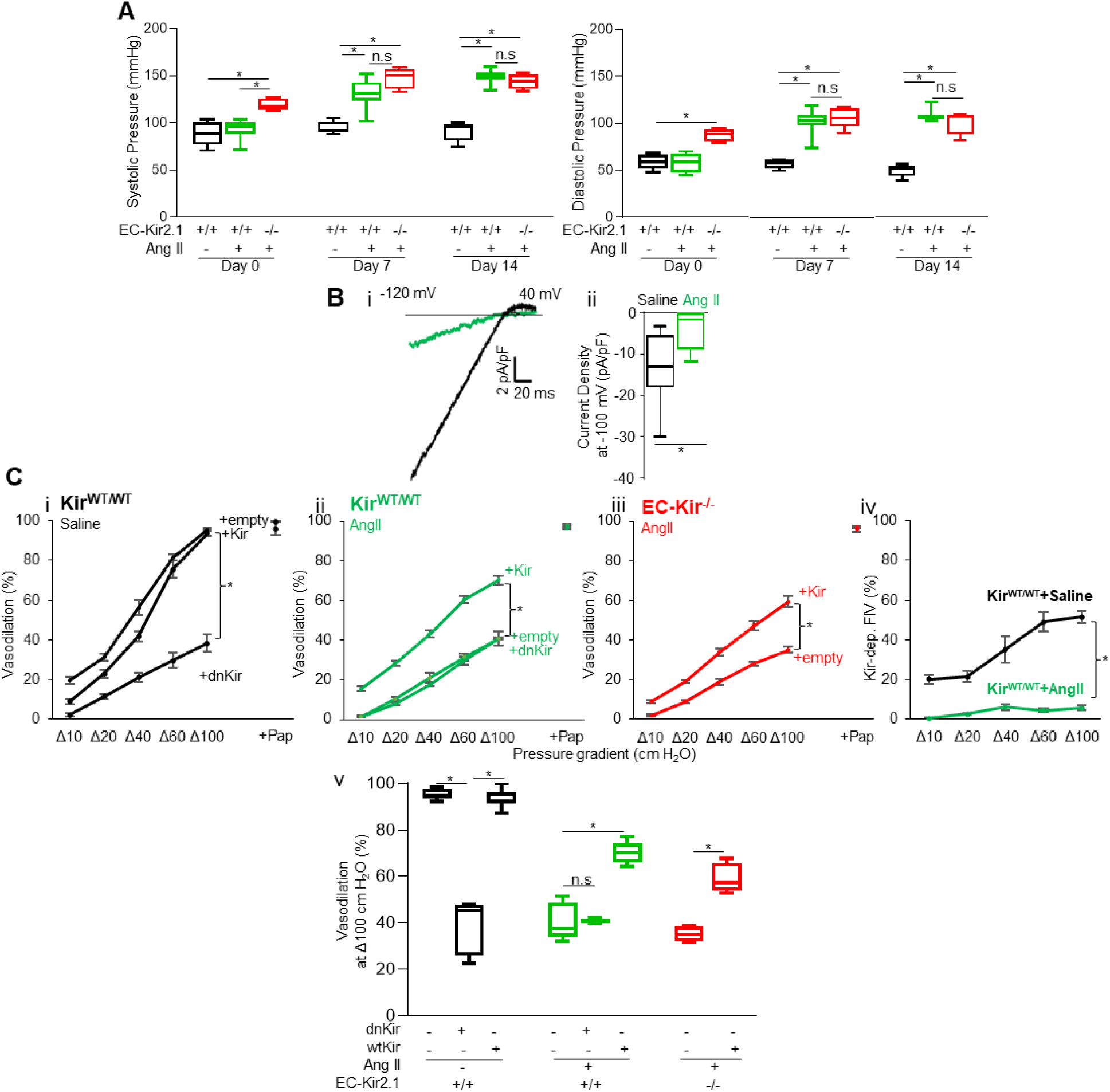
Role of endothelial Kir2.1 in AngII-induced FIV suppression. *<u>A. AngII-induced elevation of blood pressure.</u>* Systolic and diastolic pressure measured with tail-cuff in AngII-infused mice at day 0, 7, and 14. Male EC-Kir2.1^+/+^ infused with saline (black) were used as control (n=5, *p<0.05). *<u>B: AngII infusion suppresses endothelial Kir2.1</u>*. *i*. Representative currents of Kir2.1 in freshly isolated endothelial cells from mesenteric arteries of EC-Kir2.1^+/+^ mice infused with saline (black) and AngII (green). *ii*. Current density of Kir2.1 at -100 mV in freshly isolated endothelial cells from mesenteric arteries of EC-Kir2.1^+/+^ mice infused with saline (black) and AngII (green) (n=5, *p<0.05). *<u>C. AngII suppresses Kir2.1-dependent FIV</u>*. *i-iii.* FIV from mesenteric arteries in EC-Kir2.1^+/+^ (green) and EC-Kir2.1^-/-^ (red) mice infused with AngII for 14 days. EC-Kir2.1^+/+^ infused with saline (black) was used as control. Either empty Adv vector, *wtKir2.1-AdvCdh5 (Kir),* or *dnKir2.1-AdvCdh5 (dnKir)* were transfected to arteries (n=5, *p<0.05). *iv.* Kir2.1-dependent FIV (empty-dnKir) in arteries of EC-Kir2.1^+/+^ mice infused with saline (black) and AngII (green) (n=5, *p<0.05) *v.* FIV at Δ100 cm H_2_O (max flow) for all the conditions described in i-iii (n=6, *p<0.05) Pap: papaverine (100 µmol/L(µM))

**Figure 6.**
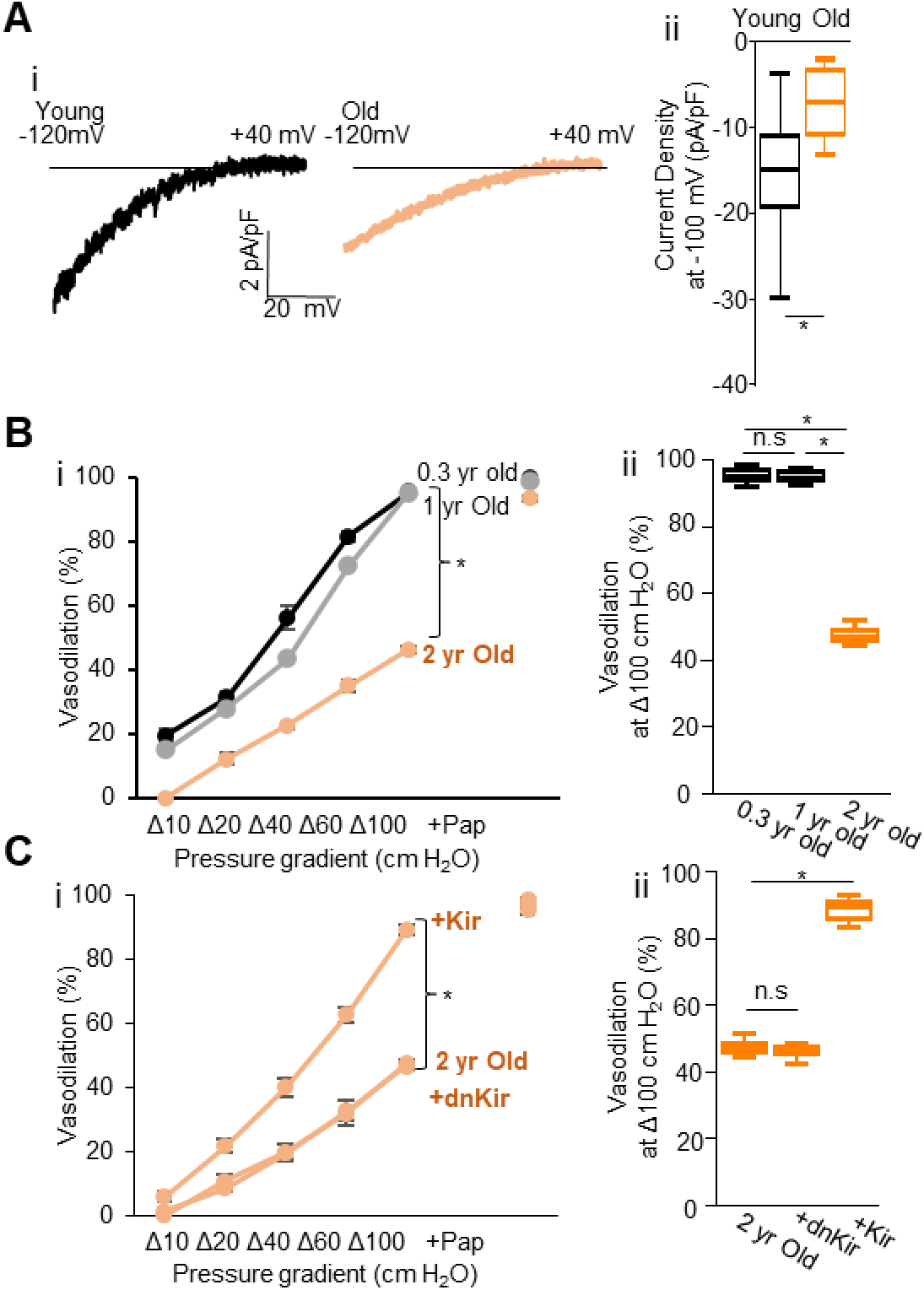
Role of endothelial Kir2.1 in FIV impairment in aged mice. *<u>A. Aging results in significant decrease in endothelial Kir2.1 currents</u>. i.* Representative Kir2.1 currents in freshly isolated endothelial cells from young (4 months old, black) and old (24 months old, orange) WT mice. *ii*. Current density of Kir2.1 at -100 mV in freshly isolated endothelial cells from young (4 months old, black) and old (24 months old, orange) WT mice (n=10 cells from 7 young mice, n=9 cells from 5 old mice). *<u>B. Suppression of FIV in aged mice.</u> i.* FIV in mesenteric arteries from 4 months (black), 12 months (grey), and 24 months (orange) old WT mice. (n=7 4-month-old mice, n=5 12-month-old mice, n=5 24-month-old mice, *p<0.05). *ii.* FIV at Δ100 cm H_2_O (max flow). (n=7 4-month-old mice, n=5 12-month-old mice, n=5 24-month-old mice, *p<0.05) *<u>C. Rescue of FIV in arteries isolated from aged mice by Kir2.1 over-expression.</u> i.* FIV in mesenteric arteries from 24 months old WT mice transfected with empty Adv vector, *wtKir2.1-AdvCdh5 (Kir),* or *dnKir2.1-AdvCdh5 (dnKir)* (n=5, *p<0.05) ii. FIV at Δ100 cm H_2_O (max flow). (n=5, *p<0.05) Pap: papaverine (100 µmol/L(µM))

The role of endothelial Kir2.1 in AngII-induced impairment of FIV is demonstrated by showing that while functional downregulation of endothelial Kir2.1 by *dnKir2.1-AdvCdh5* significantly impairs FIV of arteries isolated from EC-Kir^WT/WT^ mice infused with saline, it has no effect on the FIV of arteries from AngII-infused EC-Kir^WT/WT^ mice, indicating that AngII and loss of Kir2.1 do not have additive effects (*<u>Fig.5Ci,ii,v</u>*). Control arteries isolated from the same mice were transduced with an empty (control) Adv vector. An increase in endothelial Kir2.1 expression by *wtKir2.1-AdvCDh5* in arteries isolated from mice infused with AngII results in significant (∼75%) FIV recovery in both WT and EC-Kir2^-/-^ mice (*<u>Fig.5Cii,iii,v</u>*). The impact of AngII on the Kir2.1-dependent component of FIV is further demonstrated by subtracting the response in arteries transduced with *dnKir2.1-AdvCdh5* from the response in arteries transduced with the empty vector isolated from the same mouse (*Fig. 5Civ*). Taken together, these observations show that a significant portion of AngII-induced impairment of FIV can be attributed to the loss of endothelial Kir2.1. Since we have shown previously that there is no difference in the effects of Kir2.1 deficiency on FIV between males and females^4^, here we focused on male mice.

### Role of endothelial Kir2.1 in vascular aging

Aging is well-established as a major risk factor for hypertension, as blood pressure rises progressively with age in humans^21^. Moreover, age-related decline in FIV is well documented in both humans and animal models including mice^22^. Here, we demonstrate that a significant portion of age-related impairment of FIV in resistance arteries of advanced age mice can be attributed to a functional loss of endothelial Kir2.1 and is fully rescued by over-expression of Kir2.1 ex vivo. First, we demonstrate that endothelial cells freshly isolated from mesenteric arteries of aged mice (24 months old) exhibit significantly lower Kir2.1 current densities (∼2-fold reduction), as compared to cells isolated from young (4-month-old) mice (*<u>Fig.6Ai,ii</u>*). Consistent with the decrease in Kir2.1 currents, Kir2.1-dependent component of FIV is virtually lost in 24-months old mice: mesenteric arteries isolated from mice of this age show a significant reduction in FIV response, which is not sensitive to the functional downregulation of endothelial Kir2.1 by endothelial-specific expression of dnKir2.1 (*<u>Fig.6Bi,ii)</u>.* Furthermore, endothelial overexpression of wtKir2.1 fully rescues FIV in aged mesenteric arteries indicating that the loss of Kir2.1 functional expression in endothelial cells in advanced aging plays a critical role in age-related FIV impairment (*<u>Fig.6Ci,ii</u>*).

## Discussion

Flow-induced vasodilation (FIV) is a fundamental endothelial response that integrates multiple mechanosensitive elements to regulate vascular tone, tissue perfusion, and systemic blood pressure^1,2^. Our earlier studies identified endothelial flow-sensitive Kir2.1 channels as a key factor in flow-induced NO production and FIV. Our current findings reveal that Kir2.1 functions as key mechanistic step that links endothelial glycocalyx, to Piezo1-mediated Ca^2+^ influx and downstream activation of the PI3K/Akt1/eNOS signaling cascade. We also discovered that the loss of endothelial Kir2.1 is a key determinant of FIV impairment in AngII-induced hypertension and advanced aging.

Our first goal was to establish the mechanism by which endothelial Kir2.1 regulates flow-induced phosphorylation of Akt1, the essential step in the activation of eNOS^23^. Flow-induced Akt1 activation is known to require two coordinated steps: translocation of Akt1 to the plasma membrane and phosphorylation of Akt1 by mTORC2, which subsequently enables phosphorylation of eNOS to promote NO production and vasodilation^24^. The translocation is driven by PI3K activity, which generates PIP3 that recruits Akt1 to the membrane^14^. We present here evidence that Kir2.1 is essential for the PI3K-dependent Akt1 translocation but not for the mTORC2-dependent phosphorylation: (i) Kir2.1 deficiency abrogates flow-induced PI3K phosphorylation; (ii) overexpression of myr-Akt1, a mutant that bypasses PI3K-dependent recruitment by anchoring Akt constitutively to the membrane^25^, rescues flow-induced eNOS phosphorylation in Kir2.1-deficient endothelial cells, as well as FIV in mesenteric arteries of EC-specific Kir2.1 KO mice; (iii) downregulation of mTORC2 abrogates both flow-induced Akt1 and myr-Akt1 phosphorylation in the presence and in the absence of Kir2.1. The observation that myr-Akt1 rescues eNOS/FIV in the absence of endothelial Kir2.1 indicates that once Akt1 is recruited to the membrane, its phosphorylation is Kir2.1-independent. Also, although myr-Akt1 is considered a constitutively active form of Akt^25^, which implies that it may not be sensitive to flow, we show that myr-Akt1 is not phosphorylated constitutively under static conditions but is phosphorylated in response to flow by mTORC2. Together, our findings support a dual-input model in which Kir2.1 regulates the PI3K-dependent Akt1 membrane recruitment, while a parallel mTORC2 pathway provides the phosphorylation input.

We also discover that endothelial Kir2.1 channels are upstream from the flow activation of Piezo1 channels. Piezo channels are mechanosensitive cation ion channels that respond to physical forces, including stretch and fluid shear stress^6,7^. Our study demonstrates that expression of Kir2.1 is essential for the flow-induced Ca^2+^ influx via Piezo1: deletion of Kir2.1 abrogates flow-induced Ca^2+^ influx, despite intact functional expression of Piezo1 whereas deletion of Piezo1 does not affect Kir2.1 and its flow sensitivity. Moreover, the effect of Kir2.1 on Piezo1-mediated Ca^2+^ influx is flow specific, while Yoda1-induced Ca^2+^ influx and vasodilation are Kir2.1-independent. Moreover, prior work demonstrated that flow-induced Piezo1 activation leads to TRPV4 activation^19^. We show here that TRPV4 agonist–induced Ca^2+^ influx and TRPV4 agonist dose-dependent vasodilation are independent of Kir2.1. These results suggest that Kir2.1 is essential for the upstream mechanical activation of the Piezo1–TRPV4 signaling axis in response to shear stress.

The link between flow-activation of Piezo1 and the PECAM1/VEGFR2/VE-cadherin complex was shown to involve P2Y_2_/G_αq/11_/PLCβ signaling^38^. Briefly, flow-induced Ca^2+^ influx via Piezo1 activates ATP-permeable pannexin channels leading to ATP efflux^26^, which in turn activates P2Y_2_ receptors, engaging G_αq/11_ and stimulating PLCβ signaling. Activated PLCβ then promotes phosphorylation of PECAM1, linking upstream mechanotransduction to junctional signaling. Our observations that the loss of endothelial Kir2.1 abolished flow-induced PECAM1 phosphorylation is consistent with Kir2.1 being an upstream signaling step.

In terms of Kir2.1 channels being a primary flow sensor, as was previously suggested^27^, our earlier study discovered that this is not the case and that flow activation of Kir2.1 is abrogated by degradation of endothelial glycocalyx^5^, which is putative primary flow sensor^28,29^. Our results here identify Syndecan-1 as the specific HS proteoglycan required for Kir2.1 activation under shear stress. The significance of this finding is that it opens the possibility to interrogate the structural determinants and the mechanistic basis of Sdc1-Kir2.1 interaction that are essential for the transduction of the flow signal between the glycocalyx and the channels, which will be addressed in future studies.

Furthermore, our study demonstrates the significance of Kir2.1 in endothelial dysfunction in AngII-induced hypertension and in aging based on three lines of evidence: (i) Kir2.1 currents are significantly inhibited in both conditions, (ii) Kir2.1-dependendent FIV is virtually lost, and (iii) Kir2.1 endothelial overexpression leads to a full rescue of FIV under these pathological conditions, providing the most compelling evidence for its crucial role in FIV impairment. Mechanistically, we suggest that the functional loss of Kir2.1 in both hypertension and aging could be a result of glycocalyx deterioration. Indeed, multiple studies demonstrated that hypertension and aging are associated with deterioration (thinning, shedding, and functional loss) of the endothelial glycocalyx, caused by multiple factors^30-32^ and in particular by oxidative stress^33^. It is also well-established that degradation of endothelial glycocalyx leads to impaired response to flow resulting in a decreased availability of NO and impaired FIV^34^. Our observations suggest that Kir2.1 could be the mechanistic link between hypertension/aging-induced deterioration of glycocalyx and the loss of FIV. Moreover, as we showed previously^5^, an increase in Kir2.1 expression may allow to generate sufficient Kir current even under pathological conditions with damaged glycocalyx.

### Perspectives

Numerous studies investigated endothelial mechanotransduction and identified multiple flow signaling pathways that converge to induce FIV through the release of NO. However, understanding the mechanistic connections between different putative flow sensors, such as endothelial glycocalyx, mechanosensitive ion channels and downstream events remains a major challenge in the field. In this study, we establish a mechanistic link between glycocalyx, Kir2.1 and Piezo1 in flow mechanotransduction placing the glycocalyx at the top of a hierarchical flow-sensing cascade that proceeds through Sdc1-Kir2.1-Piezo1/TRPV4-PECAM1-PI3K/Akt1-eNOS that drives FIV, thereby defining a more unified framework for endothelial mechanotransduction. This study also identifies endothelial Kir2.1 as a critical mechanotransductive node in the loss of FIV in hypertension and aging. Furthermore, based on the capability of Kir2.1 to fully rescue the FIV response in resistance arteries isolated from hypertensive and aged mice upon overexpression, our findings suggest that this channel constitutes a new therapeutic target to improve endothelial/vascular health under such conditions.

## Supporting information

Supplemental Materials and Detailed Methods

## Abbreviations

Kir2.1: Potassium inwardly-rectifying channel 2.1
FIV: Flow-induced vasodilation
PI3K: Phosphatidylinositol 3-kinases
eNOS: Endothelial nitric oxide synthase
PECAM1: Platelet endotelial cell adhesion molecule-1
VE-Cadherin/Cdh5: Vascular endothelial cadherin
NO: Nitric oxide
AngII: Angiotensin II
MAECs: Mouse arterial endothelial cells
HAMECs: Human adipose microvascular endothelial cells
EC-Kir^-/-^ or EC-Piezo1^-/-^: Endothelial specific Kir2.1 or Piezo1 knock out
myr: Myristoylated
mTORC2: Mammalian target of rapamycin complex 2
dnKir2.1: Adenoviral vector containing Kir2.1 with dominant negative mutation
wtKir2.1: Adenoviral vector containing wild type Kir2.1
AdvCdh5: Adenoviral vector with Cdh5 promoter
TRPV4: Transient receptor potential Vanilloid 4
Sdc: Syndecan, Gpc – Glypican
LV: Left ventricle

## Novelty and Relevance

### What is new?

- We establish new mechanistic connections between different components of flow mechanotransduction
- Endothelial Kir channels are identified as essential for flow activation of other mechanosensitive ion channels
- Hypertension and aging are shown to impair endothelial Kir function

### What is relevant?

- Endothelial deficiency of Kir results in increased blood pressure in mice
- Impairment of flow-induced vasorelaxation (FIV) in AngiotensinII-induced hypertension model can be attributed to the loss of endothelial Kir
- Impairment of FIV in aging is also Kir dependent

### Clinical Pathophysiological implications

- Increase in endothelial Kir expression results in full rescue of FIV in Angiotensin-induced hypertension and in aged mice
- This indicates that endothelial Kir is a novel therapeutic target for hypertension

