## Supplemental Materials and Detailed Methods for "Flow-sensitive K^+^ channels link flow to piezo1/PI3K/Akt1 pathway"

#### Methods (detailed)

##### Animals

All mouse experiments were approved by The University of Illinois Animal Care Committee (ACC#2022137). Pathogen-free housing at the University of Illinois Animal Care Vivarium was used for all mice, which is accredited by the Association for Assessment and Accreditation of Laboratory Animal Care International (AAALAC International) and fulfilled the standards of the Animal Welfare Act, the Public Health Service Policy, and the NIH Guide for the Care and Use of Laboratory Animals. Mice were humanely treated following the institutional guidelines and performed all animal experiments in accordance with the Guide for the Care and Use of Laboratory Animals. Experiments were done to minimize the animals' pain and suffering. Animal tissues used in the experiment were extracted after euthanasia using 30% $CO_2$ followed by cervical dislocation. Cdh5.Cre$^{ERT2}$Kir2.1$^{fl/fl}$, Akt1$^{-/-}$, Cdh5.Cre$^{ERT2}$Piezo1$^{fl/fl}$, and wild type C57BL/6 mice were used for experiments. All mouse models were developed in C57BL/6 background. 5-6 months old mice were used, except specifically noted. Tamoxifen injections to induce the downregulation of endothelial Kir2.1 were performed, as described previously[1]. Tamoxifen (Sigma) was dissolved with 10 mg/ml in corn oil/ethanol (10:1) and injected 2 mg/mouse for 5 consecutive days to activate Cre enzyme. Vehicle injected litter mates were used as control. Wild type male C57BL/6 mice were maintained for 12 months and 24 months to assess the effects of aging.

##### Mouse mesenteric endothelial cell culture

Primary cultured mouse mesenteric arterial endothelial cells were isolated and maintained as previously described[2]. Mouse mesenteric arteries were extracted from 10 weeks old WT and Kir2.1$^{+/-}$ FVB male mice. Tissues were digested using Type IV collagenase (5 mg/ml, Worthington), elastase (1 mg/ml, Worthington), and the endothelial cells were sorted with anti-CD31 (PECAM1) using magnetic-activated cell sorting system (Miltenyi Biotec). Isolated mouse mesenteric arterial endothelial cells (MAECs) were cultured in EGM-2 (Lonza) media in a standard cell culture incubator. Primary cultured cells were used up to 5 passages for experimental purposes, including western blot, $Ca^{2+}$ imaging, and electrophysiological analysis, to maintain their original endothelial characteristics.

##### Human adipose microvascular endothelial cell culture

Primary cultured human adipose microvascular endothelial cells (HAMECs) were purchased from ScienCell Research Laboratories and were maintained under standard culture conditions using an endothelial cell medium and supplements (ScienCell). Low-passage cells between P2-P4 were used in all experiments. Cells were seeded at ~ 80% confluency and allowed to form a monolayer.

##### Gene silencing and qPCR

Gene silencing was performed using siRNA as previously described[3]. Primary cultured MAECs or HAMECs were incubated with commercially designed siRNAs against RICTOR, Piezo1, Sdc1, Sdc4, and Glp1 (Qiazen) using RNAiMAX Lipofectamin (Invitrogen) mixed in OptiMEM (Gibco) for 48 hours in cell culture incubator. To validate the silencing effect, primary cultured MAECs or HAMECs were lysed, and mRNA was extracted using Trizol (Invitrogen). cDNA reverse transcription was performed using High-Capacity cDNA Reverse Transcription Kit (Applied Biosystems), and qPCR was performed using Viia 7 real-time PCR system with fast-SYBR green master mix solution (Applied Bioscience). All related pre-designed primers were purchased from IDT. Data were analyzed using the $2^{-\Delta\Delta Ct}$ method. For siRICTOR and siPiezo1,

4 variants were tested, respectively. siRICTOR b and siPiezo1 b were chosen because they showed the largest reduction in mRNA expression (Supple. Fig. 1C and 2A).

**Flow-induced phosphorylation of PI3K, Akt1, eNOS, and PECAM1**

Flow-induced phosphorylation of PI3K, Akt1, eNOS and PECAM1 was measured using Western blot analysis as previously described[2]. WT and Kir2.1$^{+/-}$ MAECs were seeded in 6-well plates with 80% confluence. Cells were exposed to shear flow with 20 dyn/cm$^2$ for 3 min using cone apparatus. Cells maintained in no-flow conditions served as control. Cells were immediately rinsed with ice-cold PBS (pH 7.4) and homogenized in ice-cold RIPA buffer (Sigma) with cOmplete EDTA-free, protease inhibitor cocktail tablets (Roche), and PhosStop, phosphatase inhibitor cocktail tablets (Roche). Samples were mixed and boiled with 4x Laemmli sample buffer (Bio-Rad), supplemented with 2-mercaptoethanol (Bio-Rad). Samples were resolved by SDS–PAGE (10% acrylamide gel, Bio-Rad) and transferred onto a PVDF membrane (0.2 μm, Bio-Rad). The transferred blots were blocked with 5% non-fat milk and incubated for overnight with primary antibodies. After washing, the blots were incubated with peroxidase-conjugated secondary antibodies for 2 h and developed using iBright 1500, the ECL detection system (Thermo Scientific).

**Flow-induced vasodilation measurement**

Mesenteries were extracted from WT, Akt$^{-/-}$, EC-Kir$^{+/+}$, EC-Kir$^{-/-}$, EC-Piezo1$^{+/+}$, and EC-Piezo1$^{-/-}$ mice, as previously described[1]. Mesenteric arteries were extracted from the whole mesenteries by removing fats with forceps in HEPES buffer (pH7.4). First order mesenteric arteries between 150-200 um were cannulated to the organ chamber with glass micropipettes filled with Krebs solution (pH7.4). The inner diameter was measured after 30 min of pre-pressurization under 60 cm$H_2O$. Endothelin-1 (120 pmol/L final concentration) was used to pre-constrict up to 60% of pre-pressurized inner diameter. Flow-induced vasodilation was determined by exposing the vessels to incremental pressure gradients of Δ10, Δ20, Δ40, Δ60, and Δ100 cm$H_2O$ with maintaining the inner pressure at 60 cm$H_2O$. Inner diameter changes were measured and normalized by the following equation:

$$FIV\ (\%) = \frac{(diameters\ in\ response\ to\ flow\ -\ pre-constricted\ diameter)}{(Baseline\ diameter\ -\ pre-constricted\ diameters)} * 100$$

Papaverine (100 μmol/L) was added after the measurement of inner diameter at Δ100 cm$H_2O$ to induce endothelial-independent maximum dilation to confirm potential dilative power of vessels. To explore the role of Kir2.1 in FIV, all mouse arteries were incubated with either empty, Cdh5-promoter-driven dominant negative Kir2.1, or Cdh5-promoter-driven WT Kir2.1 expressing adenoviral vectors for 48 hours prior to measure FIV. To explore the role of myristoylated-Akt1 in FIV, mouse arteries were incubated with either empty or Cdh5-promoter-driven myr-Akt1 expressing adenoviral vector for 48 hours prior to measure FIV. GSK2193874 (100 nM, TRPV4 antagonist) was applied in the chamber and incubated for 30 min before Endothelin-1 application. Buffer solutions (in mM): HEPES – 140 NaCl, 4 KCl, 1 $MgCl_2$, 5 glucose, 10 HEPES, 2 $CaCl_2$, Krebs – 123 NaCl, 4.7 KCl, 1.2 $MgSO_4$, 2.5 $CaCl_2$, 16 $NaHCO_3$, 0.026 EDTA, 11 glucose, 1.2 $KH_2PO_4$

**$Ca^{2+}$ measurement**

$Ca^{2+}$ influx was measured using Fura-2 AM dye (Thermo Fisher) as previously described[4]. Fura-2 AM dye was dissolved in DMSO with the concentration of 300 μM right before the experiments. Afterward, Fura-2 solution was diluted in $Ca^{2+}$-free HBSS as final concentration of

3 μM. WT and Kir2.1 deficient mouse mesenteric arterial endothelial cells were cultured in either iBidi slides (flow-induced) or glass-bottom dishes (agonist-induced). Cells were washed with $Ca^{2+}$-free HBSS to remove culture media and incubated with Fura-2 solution for 20 minutes under 37 °C. Cells were washed and filled up with phenol-red free EBM2 (Lonza). Fura-2 AM fluorescence was excited at 340 and 380 nm and collected at 510 ± 80 nm using an Axiovert 100 inverted microscope (Carl Zeiss) equipped with Plan-Apo 60× with the numerical aperture (NA) 1.4 oil immersion objective, Lambda DG-4 switcher illumination system (Sutter Instruments), AxioCom Hsm camera (Zeiss), fura-2 filter set (Chroma), and AxioVision Physiology Acquisition module. Images were collected at 2-s intervals. Fluorescence ratios (F340/F380) were calculated within a circular region of interest (radius 3 μm) for each cell after subtraction of intercellular background. Data sets were normalized with signals collected before the stimulation, including flow (20 dyn/$cm^2$), YODA1 (10 μM, Piezo1 agonist), or GSK101 (100 nM, TRPV4 agonist) ($f_1/f_0$). All stimulants were applied after 30 seconds from the measurement. Flow was generated using syringe pump (Harvard). To inactivate Piezo1 and TRPV4, siPiezo1 and GSK219 (100 nM, TRPV4 antagonist) were used respectively.

#### Electrophysiology on freshly isolated endothelial cells

Freshly isolated endothelial cells from mouse mesenteries or HAMECs were used for patch clamp recording as previously described[1]. Mouse mesenteric beds were cleaned and digested using an enzyme cocktail of neutral protease (0.5 mg/ml, Worthington) and elastase (0.5 mg/ml, Worthington) in HEPES buffer for an hour at 37℃. Collagenase type IV (final concentration 0.5 mg/ml, Worthington) was added to the cocktail for 2 minutes. After the digestion, the arteries were gently dissociated on ice in the enzyme solution with syringe needles (25G). Digested arteries were pipetted several times using a glass pipette to disperse ECs into a single cell suspension. Freshly isolated cells or HAMECs were allowed to adhere on 15-mm round cover slips for patch clamp experiments. To measure flow-induced current changes, cover slips were placed in a parallel plate flow chamber as previously described[5]. Borosilicate glass pipettes (Sutter instruments) were pulled using a vertical pipette puller (Narishigue, PP-830) and had resistances between 2 and 4 MΩ. The perforated patch technique was performed by adding amphotericin B (250 μg/mL) to the pipette solution. Perforated patches were obtained within 2 to 5 minutes after formation of a GΩ seal, and accepted recordings for offline analysis maintained a series resistance between 10 and 30 MΩ. Currents from ECs were recorded using an EPC10 amplifier and accompanying acquisition and analysis software (PatchMastex, HEKA Electronik). ECs were held at −30 mV and a voltage ramp of −120 to +40 mV applied over 400 ms. Currents were low-pass filtered at 2 kHz and recordings were digitized at 10 kHz. When necessary, leak subtraction was performed offline to collect the most accurate data points at −100 mV for group analysis. To measure the flow-induced Kir2.1 currents, flow of 0.7 dyn/$cm^2$, the half maximal activation of Kir2.1 to fluid shear[6], was applied to the chamber on to the pipette sealed cells. Patch solutions were the following: Extracellular solution contained (in mM): 90 NaCL, 60 KCl, 10 HEPES, 1.5 CaCl2, 1 MgCl2, 1 EGTA, pH 7.3. Pipette solution contained (in mM): 145 KCl, 10 HEPES, 1 MgCl2, 4 ATP, 1 EGTA, pH 7.3

#### Osmotic pump implantation

Osmotic pump (2002, Alzet) implantation was performed as previously reported[7]. EC-Kir2.1$^{+/+}$ and EC-Kir2.1$^{-/-}$ mice aged 5 months were abdominally implanted with osmotic pumps (Braintree Scientific) releasing either AngII (Cayman) or saline. AngII solution in saline was prepared to be released 3 mg/kg/day and injected into the pump using 30G needle in sterile condition. Saline releasing pump was prepared as control in sterile condition. All mice were subcutaneously administered with buprenorphine (1 mg/kg) for pre-op analgesic. Mice were anaesthetized (3%)

and maintained (1-1.5%) with isoflurane during the procedure. The osmotic pumps were surgically implanted in abdominal cavity. Mice were kept for 7 days of intense monitoring during recovery.

#### Echocardiogram

Echocardiogram was recorded as previously described[2]. EC-Kir2.1$^{+/+}$ and EC-Kir2.1$^{-/-}$ mice aged 3–5 months were used for the experiments. Transthoracic echocardiography was conducted using a 40 MHz transducer (MS550D) on VisualSonics' Vevo 2100 ultrasound machine (VisualSonics, Toronto, Canada). Mice were sedated in an induction chamber using 3.5% isoflurane and then placed in the supine position on a heated stage. The heated-stage was maintained body temperature at 37°C, measured the electrocardiogram via embedded electrodes and recorded the respiration waveform via impedance pneumography. The anaesthetic plane of the mice was maintained using 1.5% isoflurane in 98.5% oxygen delivered via a nosecone. After the mice were depilated, M-mode images from the parasternal short axis view were obtained using the integrated rail system. All cardiac parameters were calculated using VisualSonics' Vevo 2100 analysis software (v. 1.6) with a cardiac measurements package.

#### Tail-cuff blood pressure measurement

Blood pressure was measured using CODA system (Kent Scientific) as previously described[2]. All mice were acclimated to restrainers daily for 7 days before the measurement. Mouse containing restrainers were placed on heating platform for 15 min to acclimate before the measurement. After 5 acclimation recordings, 15 consecutive recordings were collected to assess systolic, diastolic, mean arterial pressures, and blood flow rates. At least 10 recordings were used per experiment per mouse. The peripheral vascular resistance is defined as the ratio of mean blood pressure (mmHg) to blood flow rate (µl/min).

#### Statistics

Statistical analyses were performed using GraphPad Prizm 10. Data are presented as mean±SE. Sample size n were determined based on power analysis with 80% power and alpha 0.05. A paired or unpaired Student *t* test, 1-way ANOVA, or a 2-way with or without repeated measures ANOVA were used. Significance, in all cases, was set to $P<0.05$. Bonferroni post hoc tests were used to determine where differences existed after significance was detected with ANOVA.

### Supplemental Figure 1

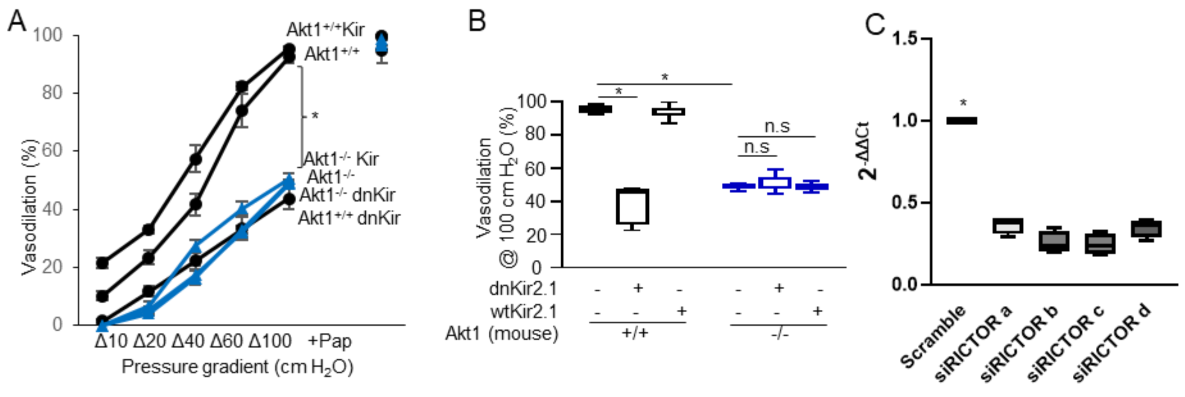


***Supplemental Figure 1.*** A. FIV measurement in mesenteric arteries from WT (Akt1$^{+/+}$, black) and Akt1$^{-/-}$ (blue) mice. Mesenteric arteries were were transfected with empty Adv vector, *wtKir2.1-AdvCdh5* (Kir), or *dnKir2.1-AdvCdh5* (dnKir). (n=5, *p<0.05) B. FIV at Δ100 cm $H_2O$ (max flow). (n=5, *p<0.05) C. mRNA expression of MAECs with scramble and 4 variants of siRICTOR. All variants showed significant reduction against Scramble. (n=4, *p<0.05)

### Supplemental Figure 2

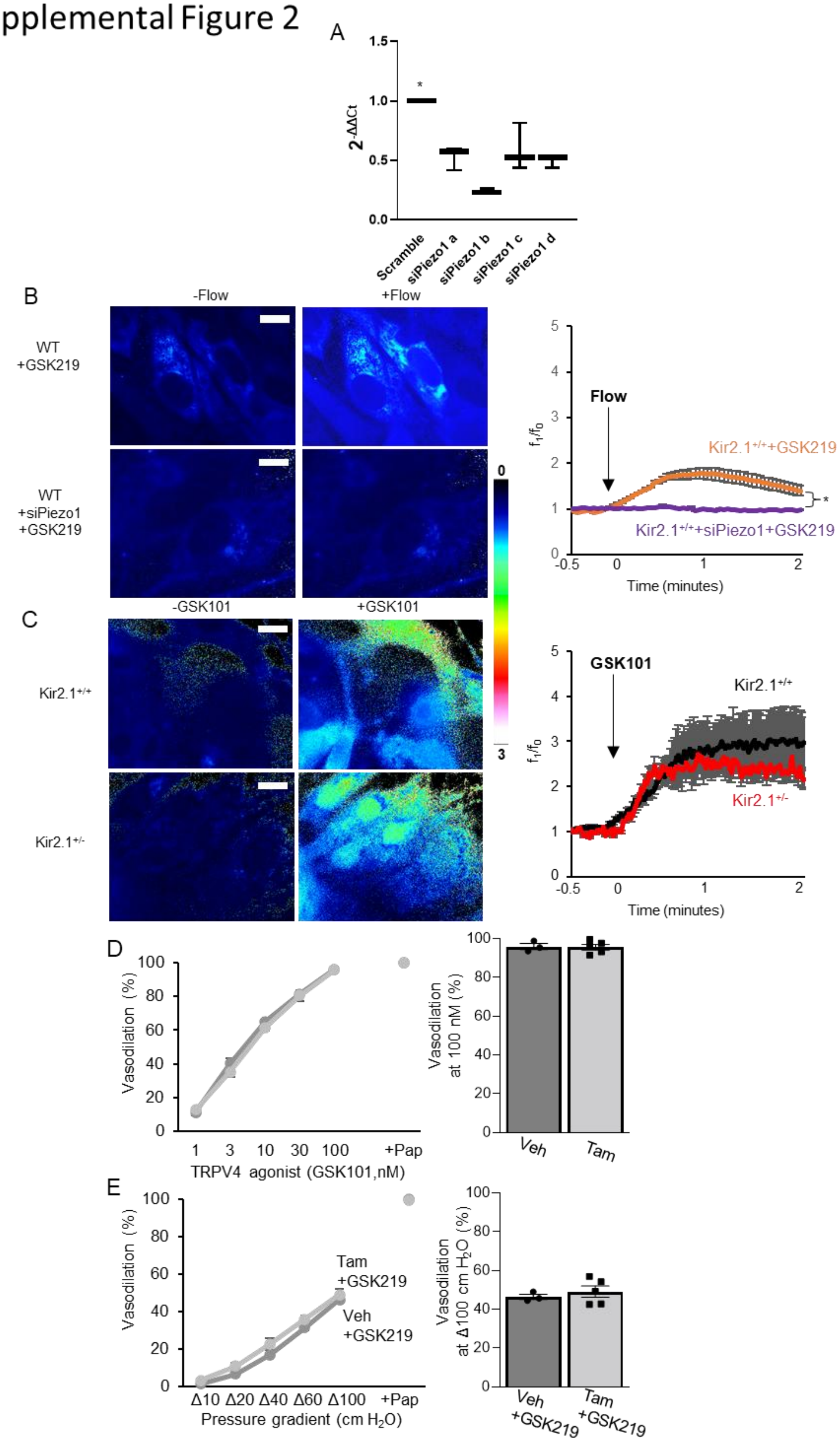

***Supplemental Figure 2.*** A. mRNA expression of MAECs with scramble and 4 variants of siPiezo1. All variants showed significant reduction against Scramble. (n=4, *p<0.05) B. (left) Representative $Ca^{2+}$ influx image before and after flow in WT MAECs applied with GSK219 or GSK219 with siPiezo1. (right) Time-dependent $Ca^{2+}$-influx curve normalized by static condition ($f_1/f_0$). (n=5 experiments, *p<0.05) C. Representative $Ca^{2+}$ influx image before and after GSK101 (100 nM) in WT and Kir2.1$^{+/-}$ MAECs. (right) Time-dependent $Ca^{2+}$-influx curve normalized by static condition ($f_1/f_0$). (n=5 experiments) D. (left) GSK101 (TRPV4 agaonist) -induced vasodilation measurement in mesenteric arteries from EC-Kir$^{+/+}$ and EC-Kir$^{-/-}$ mice. (n=5, *p<0.05) (right) Vasodilation at 100 nM (max dose). (n=5, *p<0.05) E. (left) FIV measurement in mesenteric arteries from EC-Kir$^{+/+}$ and EC-Kir$^{-/-}$ mice. (n=5, *p<0.05) (right) FIV at Δ100 cm $H_2O$ (max flow). (n=5, *p<0.05)

### Supplemental Figure 3

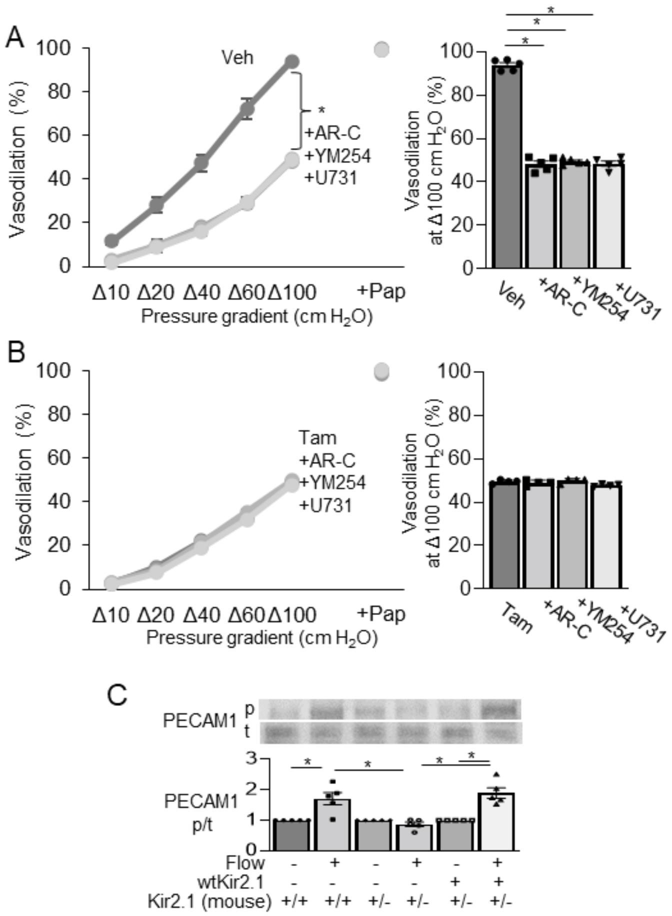


***Supplemental Figure 3.*** A, B. (left) FIV measurement in mesenteric arteries from EC-Kir$^{+/+}$ (A, Veh) and EC-Kir$^{-/-}$ (B, Tam) mice with AR-C 118925XX (10 μM, AR-C, $P2Y_2$ antagonist), YM-254890 (1 μM, YM254, $G_{\alpha q/11}$ antagonist), or U73122 (10 μM, U731, PLCβ antagonist). (n=5, *p<0.05) (right) FIV at Δ100 cm $H_2O$ (max flow). (n=5, *p<0.05) C. Western blot analysis of flow-induced PECAM1 phosphorylation in MAECs from WT and Kir2.1$^{+/-}$ mice. MAECs were transfected with empty Adv vector or *wtKir2.1-AdvCdh5*, and flow of 20 dyn/cm2 were applied for 3 min. (upper) Representative western blot, (lower) Quantitative data analysis with densitometry. (n=5, *p<0.05)

### Supplemental Figure 4

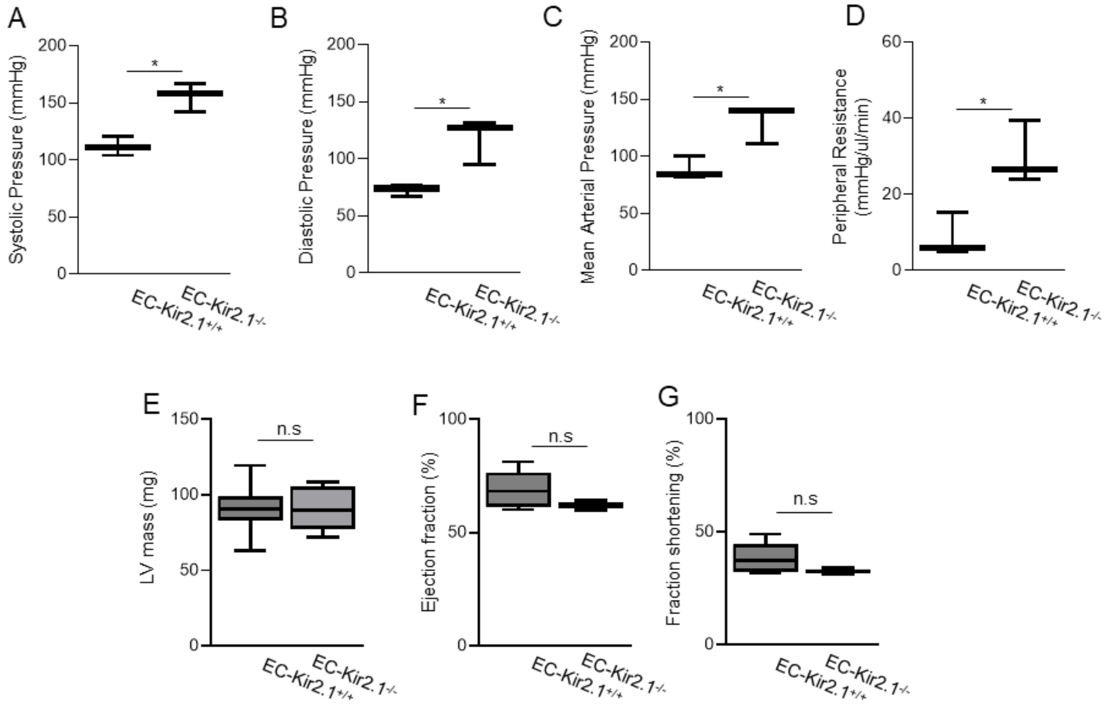


***Supplemental Figure 4.*** A. Systolic pressure, B. Diastolic pressure, C. Mean arterial pressure, and D. Peripheral resistance measured with tail-cuff from female EC-Kir$^{+/+}$ and EC-Kir$^{-/-}$ mice. (n=4, *p<0.05) E. LV mass, F. Ejection fraction, G. Fraction shortening measured in male EC-Kir$^{+/+}$ and EC-Kir$^{-/-}$ mice from echocardiogram. (n=8)
